# PseudoCell: A Multi-Valued Logical Regulatory Network to Investigate Premature Senescence Dynamics and Heterogeneity

**DOI:** 10.1101/2023.07.20.549793

**Authors:** Vinícius Pierdoná, Patrícia Lavandoski, Rafael Moura Maurmann, Guilherme Antônio Borges, Jose Carlos Merino Mombach, Fátima Theresinha Costa Rodrigues Guma, Florencia María Barbé-Tuana

**Author notes:** To whom correspondence should be addressed: Florencia María Barbé-Tuana, PhD., School of Sciences, Health and Life, Pontifical Catholic University of Rio Grande do Sul. Address: Avenida Ipiranga 6681, P12D - 401, Porto Alegre, RS, Brazil, ZIP-Code: 90619-900.

## Abstract

Premature cellular senescence is a pivotal process in aging and age-related diseases, triggered by various stressors. However, this is not a homogeneous phenotype, but a heterogeneous cellular state composed of multiple senescence programs with different compositions. Therefore, understanding the complex dynamics of senescence programs requires a systemic approach. We introduce PseudoCell, a multi-valued logical regulatory network designed to explore the molecular intricacies of premature senescence. PseudoCell integrates key senescence signaling pathways and molecular mechanisms, offering a versatile platform for investigating diverse premature senescence programs initiated by different stimuli. Validation through simulation of classical senescence programs, including oxidative stress-induced senescence (OSIS) and oncogene-induced senescence (OIS), demonstrates its ability to replicate molecular signatures consistent with empirical data. Additionally, we explore the role of CCL11, a novel senescence-associated molecule, through simulations that reveal potential pathways and mechanisms underlying CCL11-mediated senescence induction. In conclusion, PseudoCell provides a systematic approach to dissecting premature senescence programs and uncovering novel regulatory mechanisms.

## 1. INTRODUCTION

Cellular senescence is a cell state that entails the permanent cell cycle arrest associated with chronological ageing and multiple ageing-related diseases. Premature senescence can be initiated by various exogenous or endogenous stress-induced insults, such as exposure to genotoxic molecules, oxidative stress, or replication stress. This phenotype is associated with a shift in the cellular morphological and metabolic landscape, including accumulation of macromolecule damage, establishment of the senescence-associated secretory phenotype (SASP), and homeostatic disruption of metabolic pathways ^1–3^. Mechanistically, this phenotype triggers multiple, heterogeneous, and context-dependent senescence programs to achieve its cellular outcome. These molecular programs can be composed of different signaling pathways and partially depend on specific senescence-inducing stimuli ^1,4^.

In this context, understanding the diverse dynamics of premature senescence programs becomes critical. These programs, triggered by various stressors, manifest in intricate molecular and cellular responses, presenting a significant challenge in evaluating their underlying mechanisms. To address this challenge, researchers have identified different types of premature senescence programs. For instance, oncogene-induced senescence (OIS) represents a critical tumor suppressive mechanism that also acts as a barrier to tumorigenesis. The induction of OIS is primarily driven by the activation of oncogenes, such as Ras ^5^ and BRAF ^6^, or dysfunctional tumor suppressors proteins like TP53 ^7^, which promote aberrant cell survival and cellular senescence phenotype. Another important example is oxidative stress-induced senescence (OSIS). In this scenario, the escalation in oxidative stress production or the depletion of antioxidant mechanisms leads to the propagation of DNA damage and cell cycle arrest ^8^.

Considering this perspective, the systemic paradigm aims to provide a theoretical framework that can account for the inherent stochasticity and non-linearity of complex biological phenotypes ^9,10^. Within this perspective, the *in silico* study of biological networks presents itself as a powerful tool to structure and analyze the spatial and dynamic organization of these biological systems ^11–15^. Among the existing methodologies and formalisms in this field, the logical modelling of the biological networks stands out given its flexibility and high predictive power ^16–23^.

In logical modelling, each biological entity, such as genes or proteins, is represented as a node or a vertex in a graph and is associated with a discrete variable representing its activation state. The relationships between these elements are represented as links or edges in the graph. These edges can denote two types of interactions: direct interactions, such as the physical association in the formation of a molecular complex, or indirect interactions, such as the expression stimulation induced by a transcription factor. These relationships are then described in the form of logical functions. This structure allows computing the state transition of a given node in the face of changes in neighboring regulatory nodes ^22–25^. Therefore. logical modelling of biological systems is an alternative for understanding the molecular dynamics established in complex phenotypes, such as premature senescence, from a systemic perspective.

In this sense, the proposition of a logical model that encompasses the main senescent mechanisms and markers to determine the dynamics described by premature senescence programs holds significant promise for unraveling the biological complexity inherent to these processes. In this work, we present PseudoCell, a multi-valued logical regulatory network built to assist the investigation of the different dynamics in premature senescence and unfold its intrinsic heterogeneity.

## 2. MODEL SCOPE AND DEFINITION

Central to the logical modeling of regulatory networks is the initial delineation of their constituent elements. The scope of a model guides the selection of these elements, encompassing the array of regulatory pathways and biological entities essential for the modeled biological phenomena ^24^. In the context of our investigation into the dynamics underlying the establishment and perpetuation of premature senescence in humans, PseudoCell emerges as a promising regulatory network. Therefore, it becomes imperative to ascertain the involvement of pertinent pathways and biological components in this phenomenon and elucidate their interconnections.

Macromolecular damage, especially to genomic material, is an important marker in many senescence programs. Cellular insults such as oxidative stress or exposure to genotoxic materials, act as agents that potentially compromise genomic integrity ^2,3^. The accumulation of DNA damage leads to the mobilization of an extensive and complex repair machinery, collectively called DNA Damage Repair (DDR). This repair mechanism, however, is not homogeneous and different sources of damage can lead to different signaling pathways. In general, the propagation of DNA damage by genotoxic agents, such as oxidative stress, converges on the phosphorylation of the protein Tumor Protein P53 (TP53) mediated by ATM Serine/Threonine Kinase (ATM) or ATR Serine/Threonine Kinase (ATR) ^26–28^. Activation of TP53 leads to activation of cyclin-dependent kinase (CDK) inhibitors, such as Cyclin-Dependent Kinase Inhibitor 1A (p21^CIP1^) ^29^, encoded by the CDKN1A gene. The activation of proteins from the CDK inhibitors family mostly leads to cell cycle arrest through hypophosphorylation of retinoblastoma protein (pRb) ^30^.

Among the effector mechanisms of macromolecular damage, it is important to consider the role of oxidative stress. Normal reactive oxygen species (ROS) levels are important for maintaining homeostatic cellular processes. However, this system must be maintained by a delicate balance, supported by enzymatic and non-enzymatic antioxidant defenses mechanisms ^31^. The dysregulation of these mechanisms, whether through an increase in ROS production or a reduction in antioxidant mechanisms, leads to the promotion of oxidative stress, with damage to macromolecules and the expression of pro-inflammatory factors. Indeed, evidence indicates that ROS is an important factor in multiples senescence programs^32^.

Furthermore, inflammation promotes the senescent phenotype and macromolecular damage alone or in conjunction with ROS. For example, oxidative stress is responsible for increasing the production of damage-associated molecular patterns (DAMP), which act by increasing the expression of cytokines and chemokines mediated by receptors such as Advanced Glycosylation End-Product Specific Receptor (RAGE) ^33,34^. Concomitantly, evidence indicates that the expression of inflammatory mediators, such as Interleukin 1β (IL-1β) ^35^, leads to the consolidation of an oxidative phenotype. In this sense, inflammation and oxidative stress form an interdependent phenotype whose apex is the accumulation of DNA damage ^36,37^.

Furthermore, an important marker for many senescence programs is the consolidation of the SASP. SASP is characterized by the highly heterogeneous secretion of multiple soluble factors, including pro-inflammatory cytokines and chemokines (e.g. IL-1α, IL-6, and CCL2), growth factors and metalloproteinases ^38^. Evidence indicates that an important factor in the construction of this phenotype is DDR, which signals through NF- kB ^39^ to assemble most of the secretory phenotype. Therefore, this causal relationship between DNA Damage and SASP through NF-kB draws an intrinsic parallel between this phenotype and the pro-inflammatory and pro-oxidative pathways in the formation of senescence. However, using SASP as a senescence marker is limited by its high heterogeneity ^4,38^. Different senescence programs lead to the formation of distinct secretory phenotypes. Therefore, understanding how these secretory patterns relate to different senescence programs requires a systemic analysis.

In this sense, PseudoCell was modelled to comprehend pathways and molecules associated with DDR, oxidative stress, inflammation, and other auxiliary pathways, such as apoptosis and autophagy. Currently, the PseudoCell network consists of 594 nodes connected by more than 2.000 directional edges (Figure S1). A comprehensive list of the network components, their descriptions and the references are available at the PseudoCell website.

## 3. LOGICAL FORMALISM

In logical modelling, regulatory networks are depicted as a graph, *G* = (*V*, *E*), where V defines the vertices or nodes, and E represents the links or edges. Each node in this model represents a biological entity involved in the phenomenon under study, such as genes or proteins. Conversely, the links or edges abstractly represent the activation or inhibition mechanisms between two network elements ^25,40^. Also, network nodes are associated with discrete variables xi that qualitatively represent the activation level of the corresponding element ^25^. Generally, a node’s activation state represents the threshold beyond which the element initiates a specific biological function and can emulate a series of biological phenomena, including gene expression and protein phosphorylation. However, these representations are approximations of concrete phenomena designed to decrease complexity and facilitate analysis and synthesis. In multi-valued networks, such as PseudoCell, the activation state of a given node is represented by a numerical value, assuming values in the range xi = [0, Maxn], where Maxn is the maximum state value described for that component.

Finally, to each node is assigned a logical function to determine its activation state xi at a given time. The logical function, therefore, will determine the state transition of a node by combining logical and relational operators to the activation states of its regulatory nodes. The state of the logical network, at a given moment in time t, will be defined by a vector of activation states 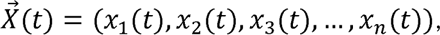 where *n* is the number of nodes in the network ^41^. However, the transition between states in a logical network depends not only on the set of logical functions that constitute it but also on the node update scheme applied. Currently, logical networks are based on two major update paradigms: synchronous and asynchronous update schemes. In the synchronous update paradigm, for each update round, all nodes have their activation state updated simultaneously, while in the asynchronous update paradigm, only one random node will be updated. PseudoCell applies a random order asynchronous update scheme, a modification to the asynchronous update scheme where, in each update round, the nodes are randomly organized into a list and then updated. The resulting dynamics can thus be evaluated throughout the system’s evolution, after a certain arbitrary period or when the model reaches one or more stable states.

## 4. PSEUDOCELL INTERFACE

PseudoCell also implements its own open-source analysis and simulation interface, written in Java, that allows users to evaluate the dynamics of the regulatory network in response to perturbations at specific nodes over time. This interface allows users to analyze PseudoCell network node behavior in different molecular contexts, such as overexpression of a protein or knockout of a gene. In this sense, through its interface, PseudoCell allows the user to define an external stimulus frequency to any element of the network (input). The stimulus frequency is measured in activation percentage (0-100%) and determines how many times that element (input) will be activated throughout the simulation.

In the PseudoCell interface, the execution time, or the number of rounds of updates that the simulation will undergo before stopping (called ticks), is defined by the user, and the simulation results are represented in the form of node activity frequency (NAF). The NAF represents the sum of the activation states that the node obtained throughout the simulation divided by the simulation time. This means that a dichotomous node (activation states ranging from 0 to 1) with an NAF of 0.87 was activated 87% of the time during the simulation. Due to the inherent stochasticity of the update scheme implemented in PseudoCell, NAF results may vary in two different simulations. This variation is important, as it brings the results of the logical model closer to its biological counterpart. To assess this variability, the user can define the number of repetitions for the simulation. Each repetition is called a sample, and the set of samples is called an experiment. This measure, therefore, allows the semi-quantitative assessment of the dynamics established by the network. It is important to note that the literature corroborates these simulation strategies, as important tools in the field, such as CellCollective, use them ^12^.

Upon completion of the experiment, the tool generates a matrix where the rows correspond to the result array of each sample (S1, S2…Sn). This matrix is then exported as a Comma Separated Values (CSV) file for user analysis. In addition, the PseudoCell interface also generates a CSV file where it reports the state of all network nodes after each update cycle. From this file, the user can determine the update cycle at which the nodes of the network entered stability or reached a steady state. This evaluation is important as it guides the choice of the minimum time for which the network should be kept running under those conditions.

The PseudoCell interface is organized into three major modules: the core module, the user interface module and the export/import module (Figure 1). Briefly, the core module is the backbone of PseudoCell, as it is responsible for computing the state transitions that drive the simulation. The user interface module is divided into three sub-modules: General, Node, and Rules Configurations. The General Configurations sub-module receives and processes the experimental conditions and simulation settings determined by the user. This includes the name of the target node, the number of times the simulation will be repeated (Sample Size, S), the number of times the network will be updated at each simulation (ticks), and the network activation stimuli.

**Figure 1.**
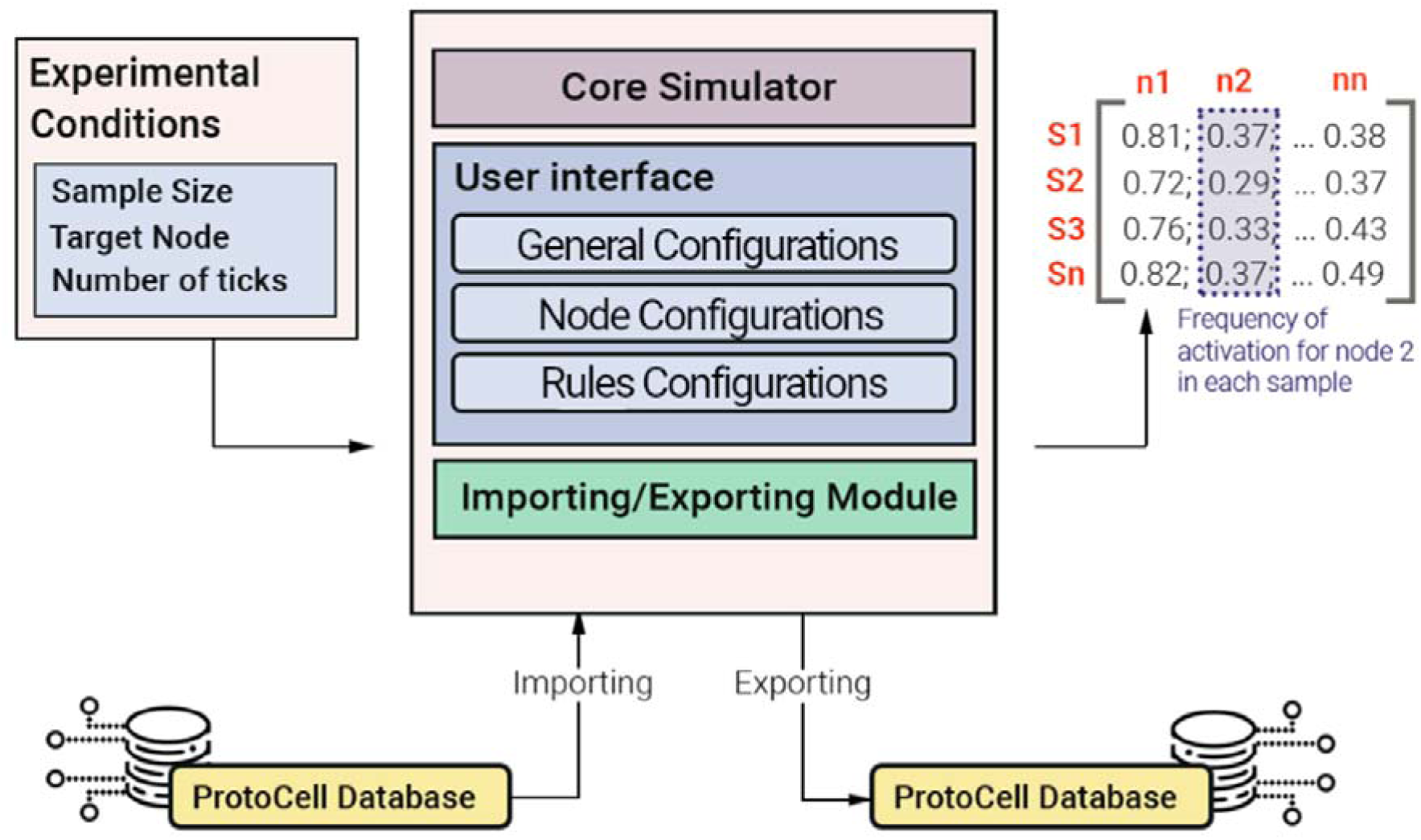
PseudoCell interface architecture. The interface is organized into three modules, User Interface, Core Simulator and Importing/Exporting Module. The User Interface module is responsible for handling the user’s experimental conditions. The Core Simulator module is responsible for carrying out the network update iterations based on the logic rules described. The Import/Export Module handles the functions for sharing the networks created by the community. The interface returns a matrix of activation frequency values as output, where the lines represent the samples (S1, S2….Sn) and the columns (n1, n2…nn) the network nodes.

We have designed PseudoCell as a primarily regulatory network for evaluating dynamics associated with senescence. However, its interface allows users to edit the original network through the Node Configurations and Rules Configurations sub-modules. These functionalities allow the original network to expand and include additional nodes. Furthermore, they also allow changing or adding logical functions to network nodes. Additionally, the user can knockout any node, which simulates the effect of biologically gene knockout or protein depletion on the network behavior. These functions allow the user to adapt PseudoCell to their experimental context.

Finally, the import/export module guarantees data persistence and allows users to share their networks with the community or absorb network updates made by the community in their experiments. Senescence is a vast and constantly growing field of study, so it is essential to allow users to add nodes, change the network and share their updates with the community.

## 5. Validation of PseudoCell Through Simulation of Classical Senescence Programs

To assess the accuracy of PseudoCell, we conducted simulations replicating classical senescence programs. Specifically, we emulated oxidative stress-induced senescence (OSIS) by perturbing the node related to CYBB at a frequency of 50% and oncogene-induced senescence (OIS) by constitutive expression of AKT1 and Ras nodes. All simulations were maintained for 1.000 ticks, ensuring that most nodes reached a steady state by the end of the simulation, and were repeated 30 times (n=30). The network’s NAF were expressed as a heatmap where the rows represent the nodes, and the columns represent the experimental groups. Subsequently, we compered the nodes modulated by each of the stimuli with empirical data available in literature. To determine which nodes had their activities modulated by the provided perturbations, we verified the differential activation of the nodes using R language and the clusterProfiler ^42,43^ and limma packages^44^.

### 5.1. Validation of Oxidative Stress-Induced Senescence Molecular Signature

In the oxidative stress-induced senescence (OSIS) model, the stimulation of the CYBB node resulted in increased activity of proteins associated with DDR and cell cycle arrest, such as H2AX, ATM, MDC1, TP53BP1, TP53 and p21^CIP1^ (Figure 2A-C and Supplementary Table 1S). Interestingly, these findings are in line with the molecular markers commonly found in cellular senescence induced under pro-oxidative conditions. While low to moderate levels of ROS serve as essential signaling molecules involved in cellular homeostasis and adaptive responses, excessive ROS accumulation can induce oxidative damage to biomolecules, such as DNA, proteins, and lipids, ultimately leading to cellular dysfunction and senescence. Accumulation of oxidative stress-derived DNA damage trigger DDR signaling, phosphorylating H2A Histone Family Member X (γH2AX), while also activating ATM, TP53BP1, and MDC1 ^32,45,46^. As in our model, DDR signaling converges on the phosphorylation of TP53, particularly at Ser15 and Ser20, leading to p21^CIP1^ expression and cell cycle arrest ^47,48^. Moreover, evidence indicate that ROS-mediated activation of p16^INK4a^ and inhibition of CDK4 e 6, are also important in consolidation of the senescent phenotype under oxidative stress ^49,50^. These elements were also found to respond to *in silico* NADPH Oxidase 2 (CYBB) stimulation in our model (Figure 2D). Lastly, we observed the expression of SASP-associated markers, including NF-kB, a significant transcription factor for stress-induced senescence and a key player responsible for the secretion of IL-6 and IL-8, two major constituents of the SASP ^51^ (Figure 2E). These *in silico* results supports the idea that our model is capable of producing a molecular activation profile mirroring the response seen in living organisms after exposure to an oxidative stressor.

**Figure 2.**
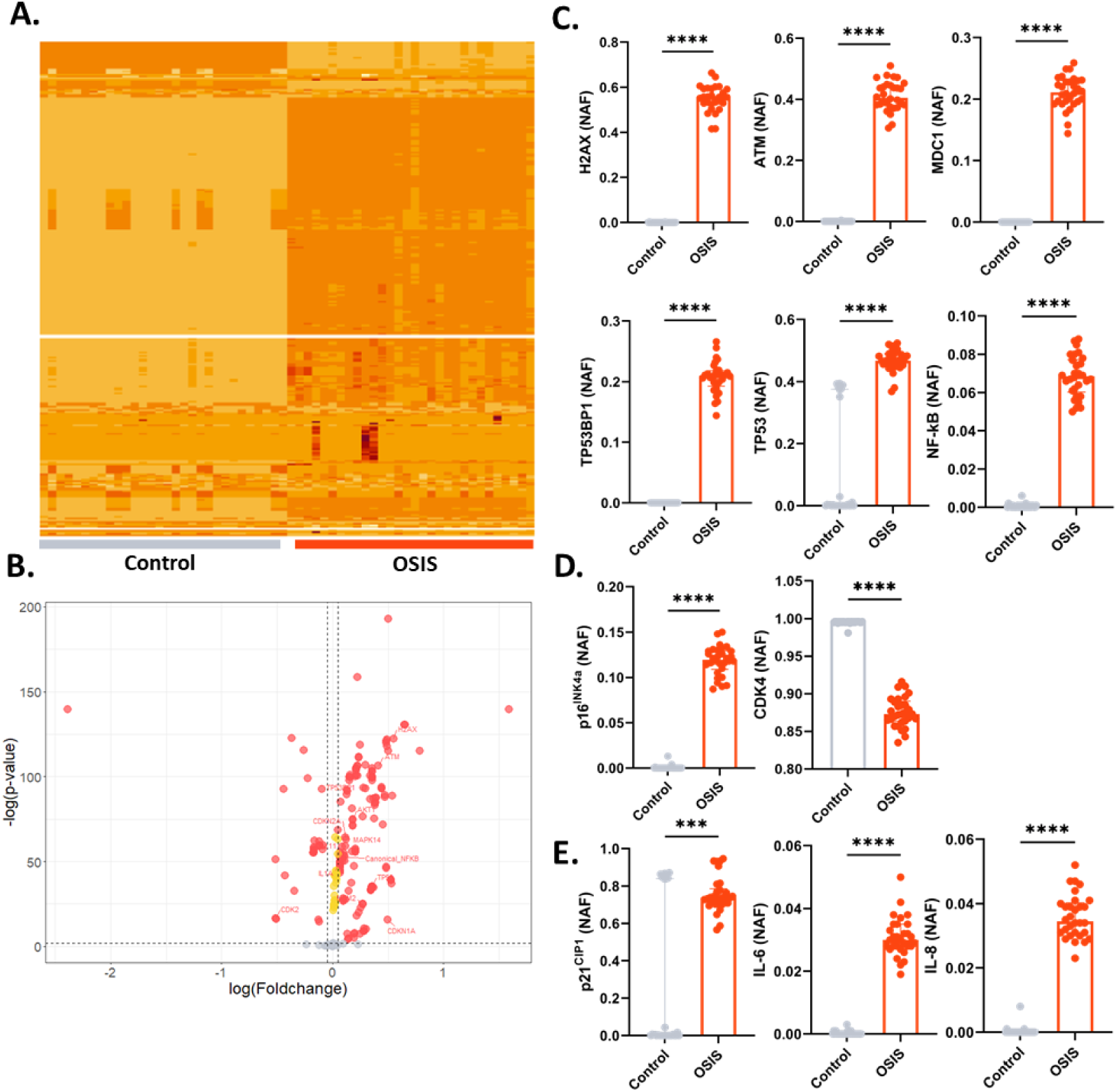
*In silico* stimulation of OSIS inducing factor CYBB leads to activation of molecular elements similar to those expected in biological models. **A.** Heatmap demonstrating the systemic response of the network to 50% frequency stimulus of the CYBB node. In the graph, the lines represent the network nodes, while the columns represent the samples from both experimental groups. **B.** Volcano plot representing differential activation nodes in OSIS group compared to the control. Red circles indicate nodes with a p-value < 0.01 and a fold change greater than 0.05. Yellow circles represent nodes with only a p-value < 0.01 and were not considered differentially activated. Gray circles represent no differentially activated nodes. **C.** OSIS leads to the expression of DNA damage and cell cycle arrest markers. **D.** OSIS leads to increase in p16 activation and decrease in CDK4. **E.** OSIS leads to SASP factors expression. Data are presented as median and IQR. Significant differences considered P<0.001 (***) and P<0.0001 (****). Normality data distribution was assessed using the Shapiro-Wilk normality test. Mann- Whitney U-test was used for non-parametric samples. NAF: Node Activation Frequency.

### 5.2. Validation of Oncogene-Induced Senescence Molecular Signature

Furthermore, we also assessed the network’s response to the overexpression of two senescence-associated oncogenes, Ras and AKT1. Oncogene-induced senescence (OIS) represents a critical tumor suppressive mechanism that acts as a barrier to tumorigenesis. The induction of OIS is primarily driven by the activation of oncogenes, such as Ras, BRAF, and Myc, which promote aberrant cell survival and cellular senescence phenotype. Similarly to stress-induced senescence, upon Ras superexpression (OIS-Ras), the cell undergoes DNA damage and oxidative stress pathways activation, leading to upregulation of TP53, p16^INK4a,^ and SASP factors ^5^. The p16^INK4a^ tumor suppressor pathway acts as a critical regulator of OIS by inhibiting the activity of CDK4 and CDK6, thereby preventing the phosphorylation of the retinoblastoma protein (Rb) and subsequent cell cycle progression. In our model, we also observed the modulation of these markers mediated by the constitutive expression of Ras (Figure 3A-B and Supplementary Table 2S).

**Figure 3.**
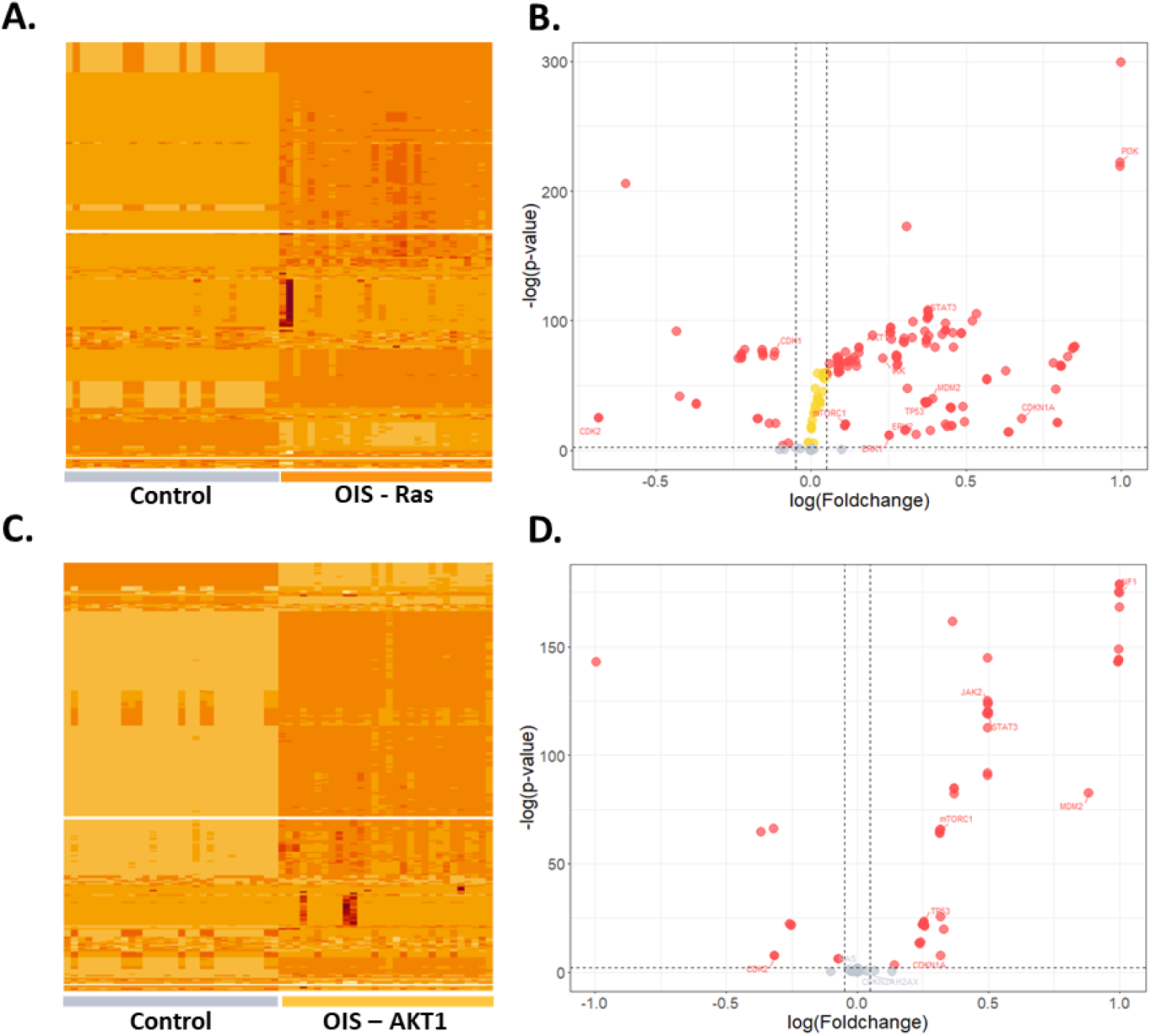
*In silico* stimulation of OIS Ras and AKT1-dependent leads to activation of molecular elements similar to empirical evidences. **A.** Heatmap demonstrating the systemic response of the network to constitutive Ras activation. In the graph, the lines represent the network nodes, while the columns represent the samples from both experimental groups. **B.** Volcano plot representing differential activation nodes in Ras stimulated group compared to the control. Red circles indicate nodes with a p-value < 0.01 and a fold change greater than 0.05. Yellow circles represent nodes with only a p- value < 0.01 and were not considered differentially activated. Gray circles represent no differentially activated nodes. **C.** Heatmap demonstrating the systemic response of the network to constitutive AKT1 activation. In the graph, the lines represent the network nodes, while the columns represent the samples from both experimental groups. **D.** Volcano plot representing differential activation nodes in AKT1 stimulated group compared to the control. Red circles indicate nodes with a p-value < 0.01 and a fold change greater than 0.05. Gray circles represent no differentially activated nodes.

Contrastingly, in the AKT-induced senescence (OIS-AKT1) model, the overexpression of the oncogene AKT1 leads to the consolidation of senescence through a distinct phenotypic pattern. Significantly, AKT1 overexpression contrasts with Ras-dependent OIS and OSIS by not involving the activation of pathways associated with DNA damage. Evidence indicates that AKT1 overexpression induces senescence through TP53 mTORC1-dependent activation and reduction of MDM2 activity in the absence of DNA damage ^52,53^. Moreover, in contrast to other models, cell cycle arrest in OIS AKT1 mediated appears to be dependent on p21^CIP1^, with no involvement of p16^INK4a^. Interestingly, our model was able to replicate these differences found among different senescence programs, indicating activation of the PI3K/AKT/mTORC1 pathway and markers such as TP53 and p21 without activation of DNA damage markers like H2AX and p16 (Figure 3C-D and Supplementary Table 3S). Also, findings suggest that OIS AKT1-induced leads to NF1-dependent suppression of Ras signaling^54^.

### 5.3. Senescence inducing stimuli leads to G1 cell cycle arrest *in silico*

Additionally, most senescence-inducing stimuli tested in our model led to an increase in G1 node activity, associated with reduced mitosis (Figure 4A). This is an important finding as many senescence models indicate G1 cell cycle arrest. For example, human fibroblasts subjected to gamma radiation-induced senescence exhibited G1 cell cycle arrest dependent on DNA damage and prolonged activation of TP53 and p21^CIP1^ ^55^. Furthermore, the cell cycle arrest in G1 positively correlates with the activity of the p21^CIP1^ and TP53 nodes in our model, replicating the expected behavior in a biological system (Figure 4B).

**Figure 4.**
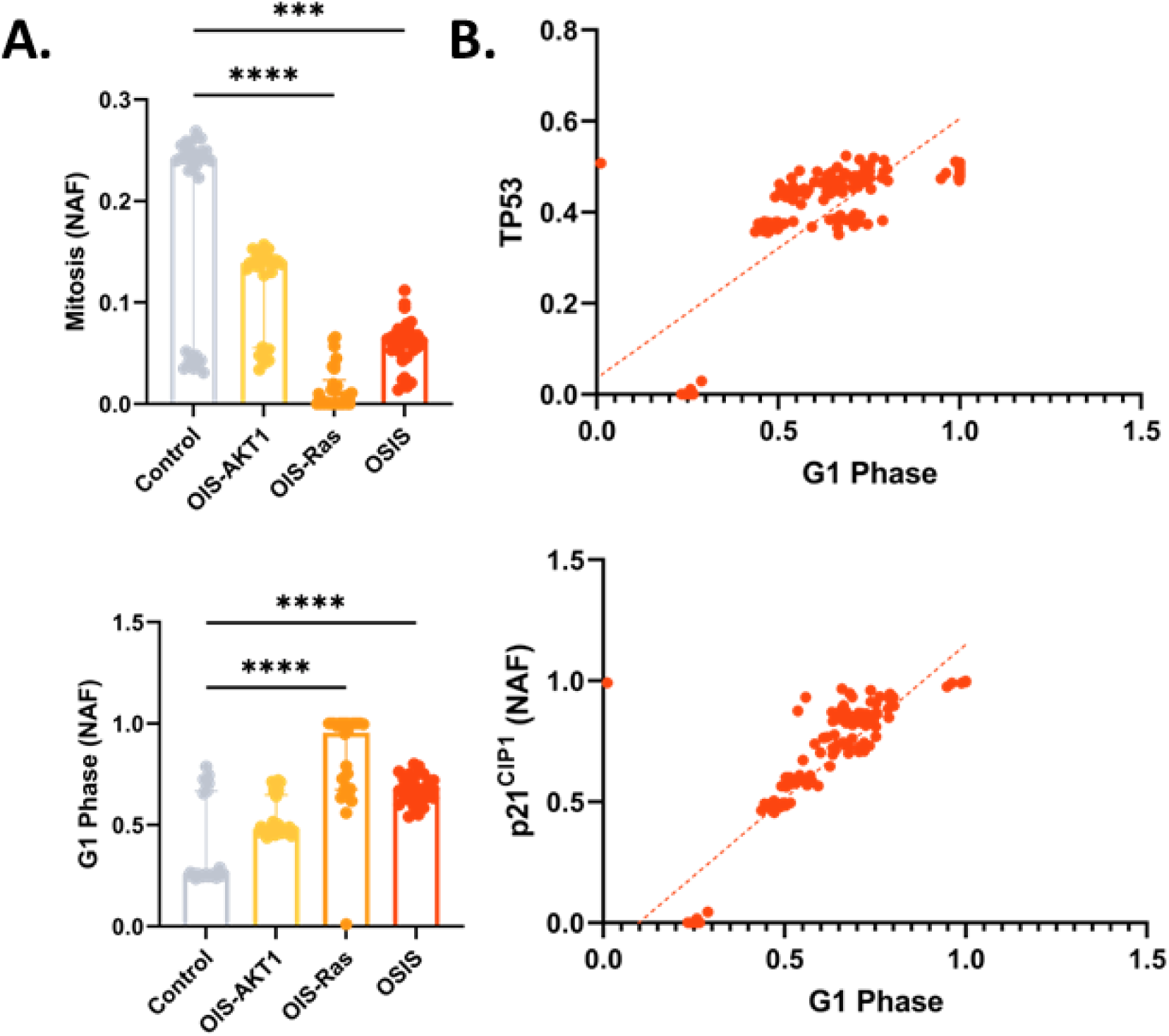
*In silico* cell cycle arrest simulation in different premature senescent models. **A.** Increased activation of the G1 Phase node in most senescence-inducing stimuli and reduced mitosis. Data distribution normality assessed by Shapiro-Wilk test. Intergroup differences assessed by Kruskal-Wallis test followed by comparation of each group’s mean rank with the rank of control group. The data presented are expressed as the median and Interquartile Range. **B.** Positive correlation between G1 cell cycle arrest and TP53 or p21 activity. Significant differences considered when P<0.001 (***) or P<0.0001 (****). NAF: Node Activation Frequency.

Collectively, these findings demonstrate that PseudoCell is capable of satisfactorily emulating different programs of OSIS and OIS. In particular, we have shown that this regulatory network can delineate specific markers for certain stimuli and the molecular peculiarities evoked by them, thereby providing insights into the molecular and cellular mechanisms underlying each senescence program.

## 6. Validation: Exploring the Role of CCL11 in Cellular Senescence

Recent literature suggests that CCL11 act as a premature aging factor and was shown to be increased in elderly^56^. Studies have demonstrated that, in young mice, the serum increase of this chemokine impairs neurogenesis, learning, and memory associated with ageing ^56^. This relationship is central in pathological conditions associated with ageing, such as severe or treatment-refractory asthma. In these individuals, as opposed to treatment-responsive cases, levels of CCL11 circulating in peripheral blood are negatively correlated with telomere length, an important marker of cellular ageing ^57^. Furthermore, evidence corroborates the association of CCL11 with ageing and senescence by demonstrating the age-dependent increase of this molecule in human peripheral blood^58^. Despite this, little is known about the molecular and intracellular events that explain the association of this chemokine with cellular ageing.

To validate the PseudoCell regulatory network’s capacity to assist in the study of novel molecules in premature senescence induction, we sought to evaluate molecular aspects associated with increasing concentrations of CCL11. For this purpose, PseudoCell was stimulated with perturbations of the CCL11 node at increasing frequencies: 100% (constitutive expression), 50% (high expression), 25% (medium expression), 12.5% (low expression), and 0% (control). The network was then updated for 1.000 ticks, and the simulations were repeated 30 times (n=30) for each experimental group.

Initially, we sought to verify if the *in silico* stimulation with CCL11 was able to replicate empirical results found in the literature. For that, we assessed the activation frequency of nodes associated with the PI3K/AKT and MAPK14 pathways. In this experiment, we verified frequency-dependent increase in PI3K, AKT1, and MAPK14 nodes (Figure 5A). These results align with the literature, where an increase in the expression or phosphorylation of these proteins was observed after stimulation with CCL11, evidencing the accuracy of our model in emulating the CCL11 biological function *in silico* ^59–62^. Furthermore, corroborating its role as potential inducer of senescence, we also verified the expression of classical markers of multiple senescence programs such as TP53, p21^CIP1^, and p16^INK4a^ (Figure 5B). Being able to replicate the ageing phenotype promoted by CCL11, we sought to understand which mechanisms could explain this phenomenon.

**Figure 5.**
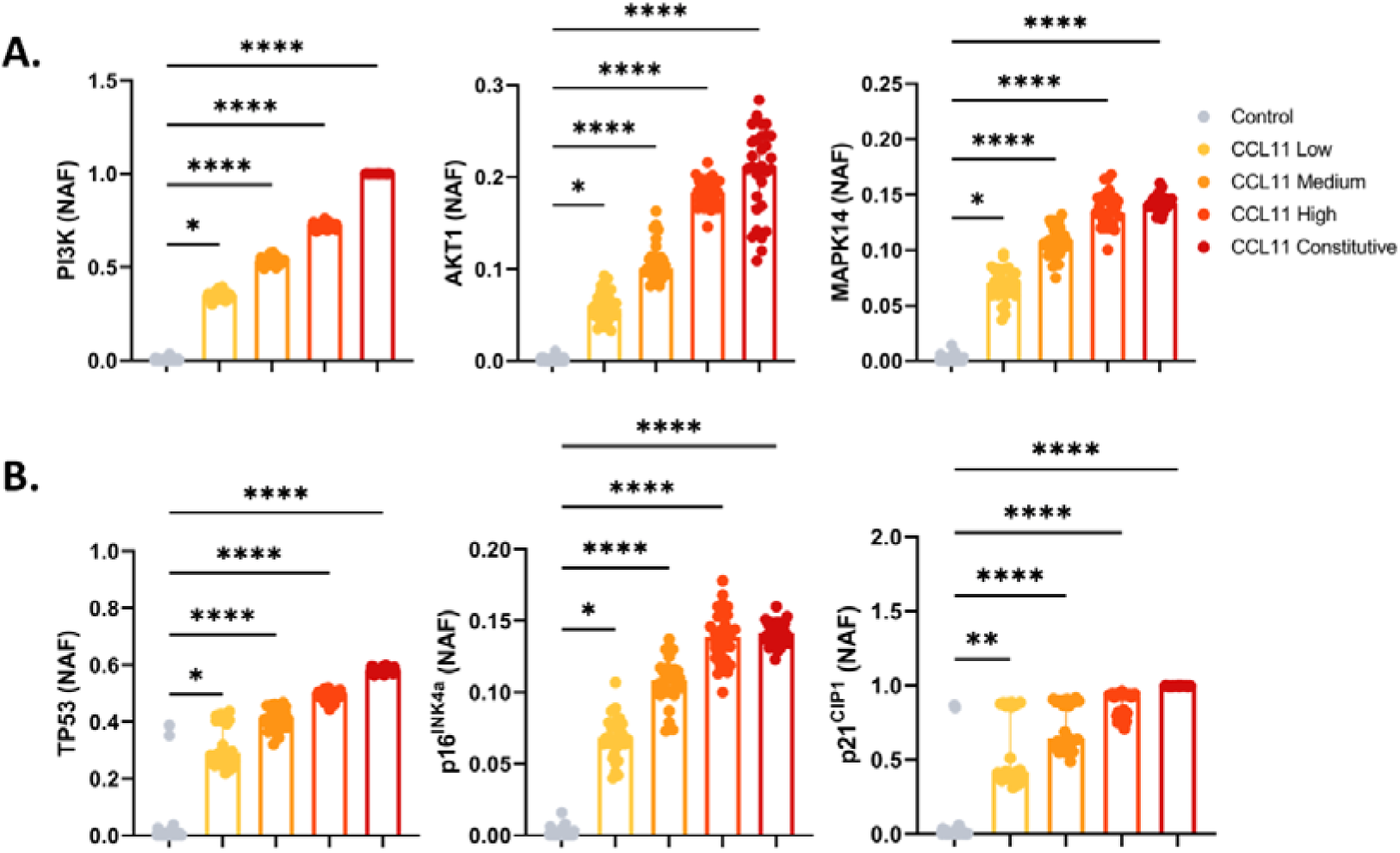
CCL11 induces activation of nodes associated with PI3K/AKT, MAPK14 and cell cycle arrest pathways in PseudoCell’s network. **A.** The PI3K, AKT1, and MAPK14 nodes had their activity modulated in a frequency-dependent manner when disturbed by CCL11. **B.** Activation of TP53, p16 and p21 node after CCL11 stimulation. Data distribution normality assessed by Shapiro-Wilk test. Intergroup differences assessed by Kruskal-Wallis test followed by comparation of each group’s mean rank with the rank of control group. Significant differences considered when P<0.05 (*), P<0.01 (**), and P<0.0001 (****). NAF: Node Activation Frequency.

Therefore, we assess the comprehensive molecular signature promoted by stimulation at increasing frequencies with CCL11 and investigate the possible mechanisms responsible for CCL11-mediated senescence induction (Figure 6A and Supplementary Table 4S). To assess consequences of CCL11 stimulation, we investigate the differentially activated nodes in the experimental groups compared to the control (Figure 6B). Sequentially, we evaluated the signaling pathways and biological functions enriched in this list of genes and proteins. Our results show an important prevalence of pathways associated with DNA damage and repair (Figure 6C and Supplementary Table 5S). This finding is expressed by an increased frequency of activation of nodes such as Double Strand Break (DSB), ATM, and H2A Histone Family Member X (H2AX). Furthermore, there was an increase in the activation of nodes associated with the ROS production pathway, such as the Superoxide node (O2^-.^) and the genes that codify for NADPH Oxidase 2 (CYBB) and Rac Family Small GTPase 1 (RAC1) (Figure 6D). Furthermore, these results were validated in peripheral blood mononuclear cells (PBMC) in vitro (Figure 2S).

**Figure 6.**
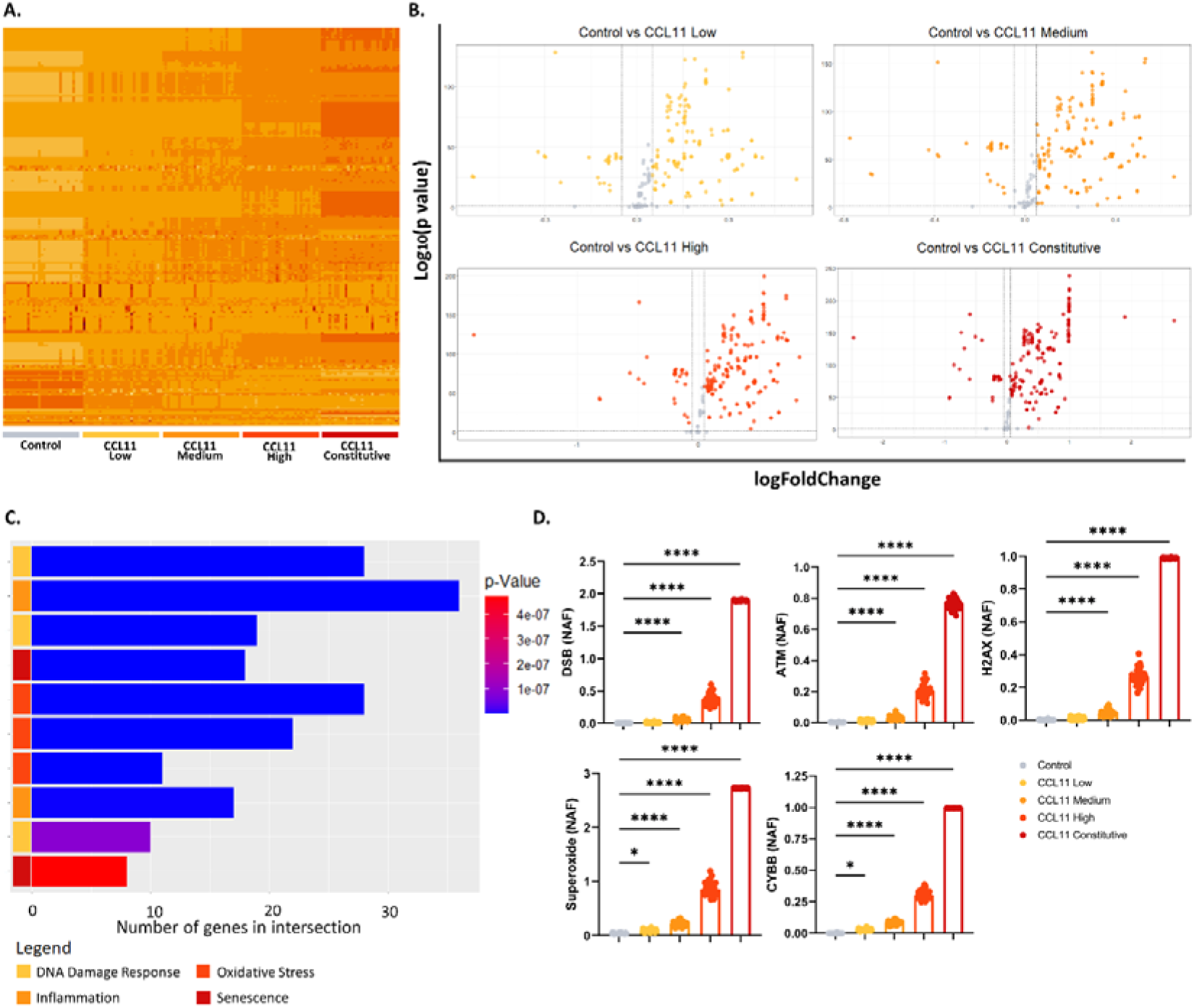
*In silico* evaluation of increasing frequency of CCL11 node activation. **A.** Heatmap representing alteration in activation frequencies patterns for Control, CCL11 Low, CCL11 Medium, CCL11 High and CCL11 Constitutive groups. **B.** Vulcano plot showing the differentially activated nodes in Controls in contrast to CCL11 Low, Medium, and Constitutive (p-Value<0.01 and |FC|>0.05). Negative FC values were considered downregulated, while positive values were considered upregulated. **C.** Top enriched pathways in CCL11 High. **D.** CCL11 activation frequencies induce increase in DSB, ATM, H2AX, Superoxide and CYBB nodes activation (p<0.0001). Data distribution normality assessed by Shapiro-Wilk test. Intergroup differences assessed by Kruskal- Wallis test followed by comparation of each group’s mean rank with the rank of control group. Significant differences considered when P<0.05 (*), P<0.01 (**), and P<0.0001 (****). NAF: Node Activation Frequency.

To verify the causal relationship between ROS production in CCL11 pathway activation and the promotion of DNA damage, the system was challenged with increasing frequencies of CCL11 node activation associated with CYBB node knockout. The results were expressed in a heatmap (Figure 7A). In this analysis, CYBB knockout, which encodes for the protein gp-91phox, the beta subunit of the NADPH oxidase enzyme necessary for the formation of ROS through the activation of the CCL11 pathway, inhibited the activation of nodes associated with the DNA damage response (DSB, ATM, and H2AX) (Figure 7B). Furthermore, inhibition of oxidative stress CCL11-induced production also led to reduced activation of TP53, but not p21^CIP1^ (Figure 7C). This indicates that ROS production has a partial role in the induction of senescence mediated by CCL11. These findings indicate that CCL11 induce a senescent phenotype with involvement of ROS formation and DNA damage, similar to that induced by cellular stress. In this sense, CCL11 emerges as a potential inducer molecule of senescence. Furthermore, our network was able to identify possible molecular mechanisms by which it could lead to this phenotype and list probable markers for its analysis.

**Figure 7.**
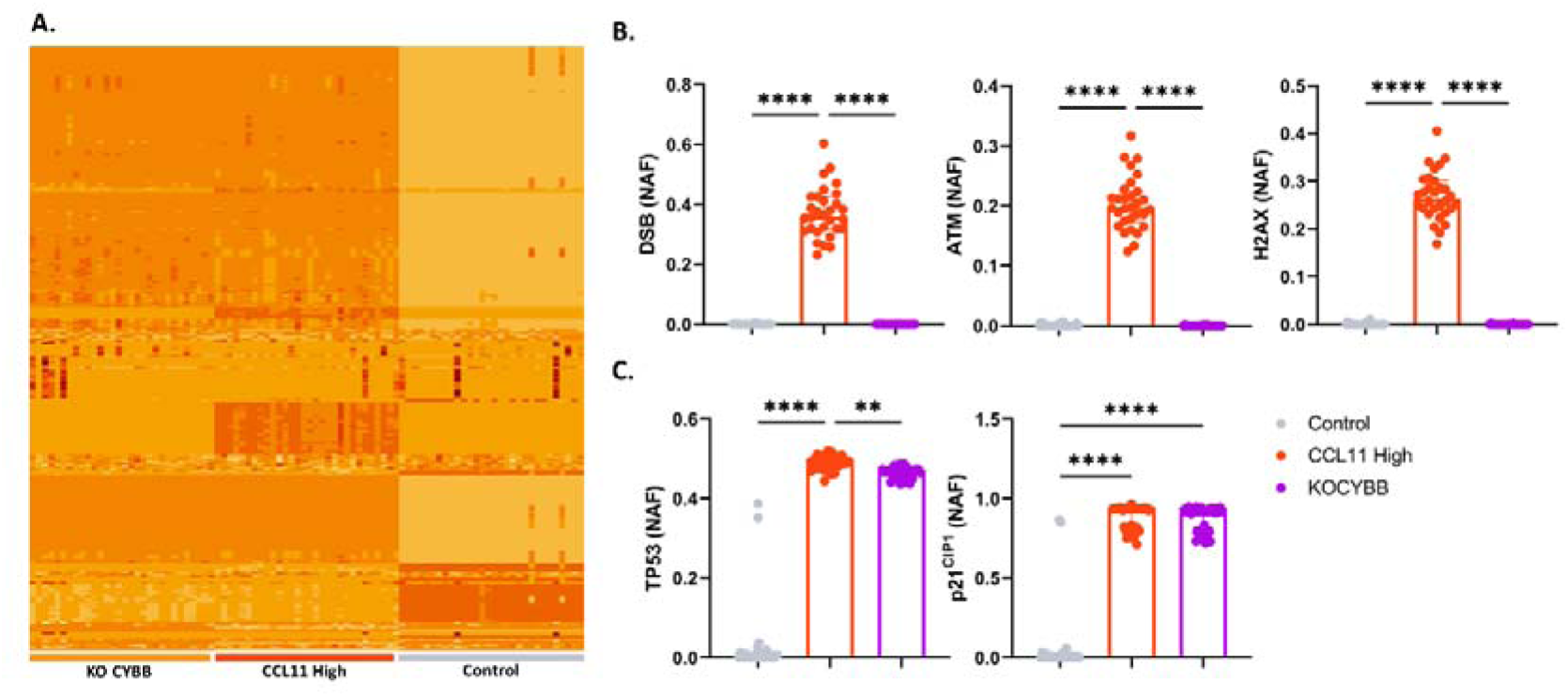
Effects of CYBB node suppression on DNA damage response and senescence markers. **A.** Heatmap representing alteration in activation frequencies patterns for Control, CCL11 High and CYBB Knockout (KO CYBB) groups. **B.** CYBB knockout suppresses activation of nodes associated with DNA damage, DSB, ATM and H2AX. **C.** CYBB knockout reduces TP53 activation, but not p21. Data are presented as median and IQR. Data distribution normality assessed by Shapiro-Wilk test. Intergroup differences assessed by Kruskal-Wallis test. Significant differences considered P<0.01 (***), and P<0.0001 (****). NAF: Node Activation Frequency.

### 6.4. CONCLUSION

In this study, we introduced PseudoCell, a multi-valued logical regulatory network designed to unravel the intricate dynamics underlying cellular senescence. By comprehensively integrating key signaling pathways and molecular mechanisms associated with senescence, PseudoCell offers a versatile platform for investigating the diverse senescence programs initiated by various cellular stimuli. Our analysis demonstrated that PseudoCell accurately recapitulates classical senescence programs, including oxidative stress-induced senescence (OSIS) and oncogene-induced senescence (OIS), by replicating molecular signatures consistent with empirical observations. Furthermore, we validated the utility of PseudoCell in exploring the role of novel molecules in cellular senescence, exemplified by our investigation into the effects of CCL11 stimulation. By simulating varying concentrations of CCL11 and analyzing the resultant molecular signatures, we elucidated potential pathways and mechanisms contributing to CCL11-mediated senescence induction.

It is important to note that PseudoCell does not claim to encompass all mechanisms associated with senescence in its current version. For instance, telomeric dynamics, a crucial aspect of senescence regulation, are not explicitly included in our model. However, future iterations of PseudoCell can expand its scope to incorporate additional components and refine existing interactions, thereby enhancing its comprehensiveness and accuracy.

In conclusion, PseudoCell emerges as a resource for investigating the complex dynamics of cellular senescence, offering a systematic approach to dissecting premature senescence programs, and uncovering novel regulatory mechanisms.

---

**Figure 1S.**
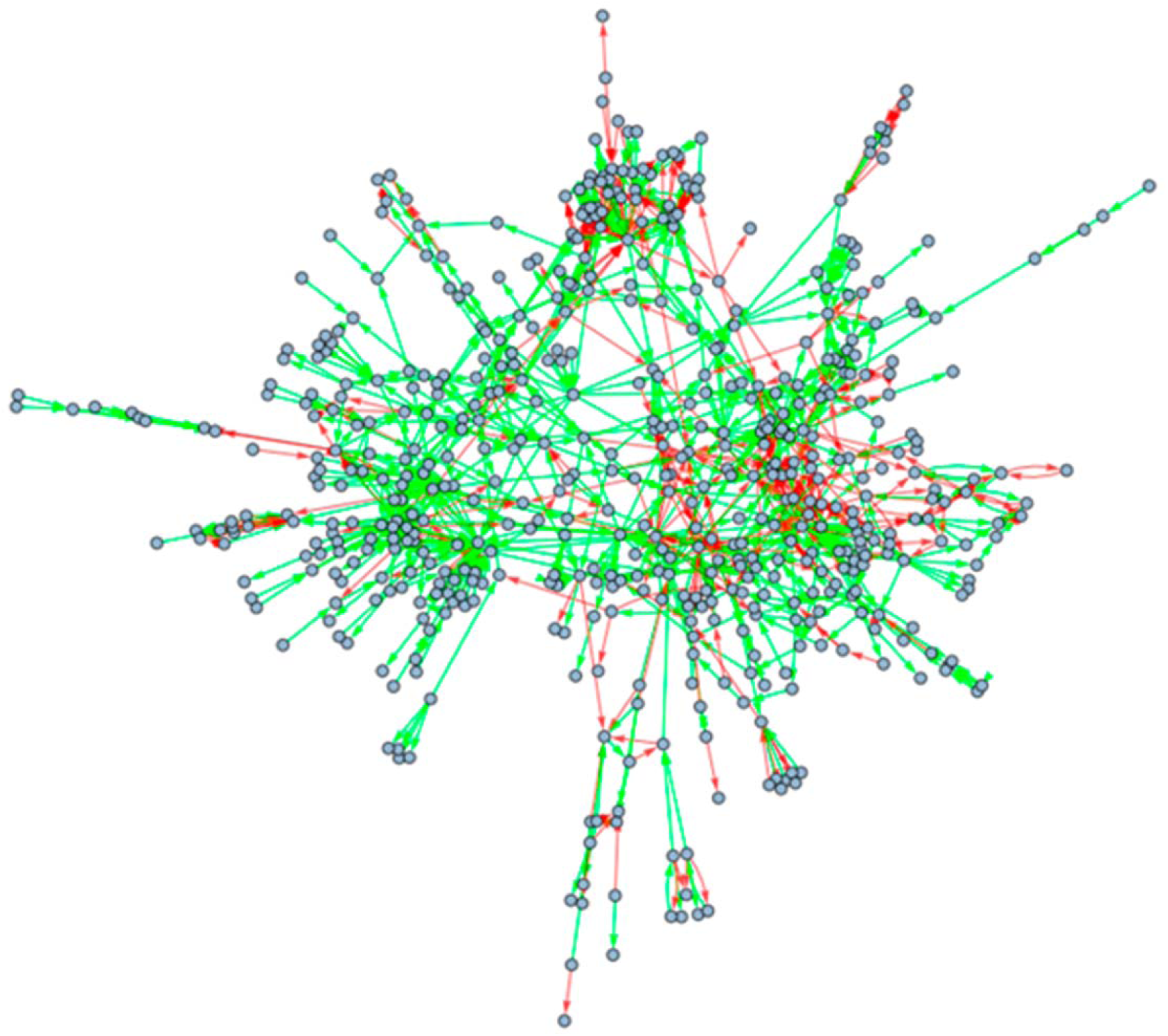
Topological structure of PseudoCell’s network. Interaction graph built from the logical model, where proteins, genes, metabolites and other biological elements are represented by circular nodes. Green edges represent the positive connections, while the red ones represent the negative connections.

**Figure 2S.**
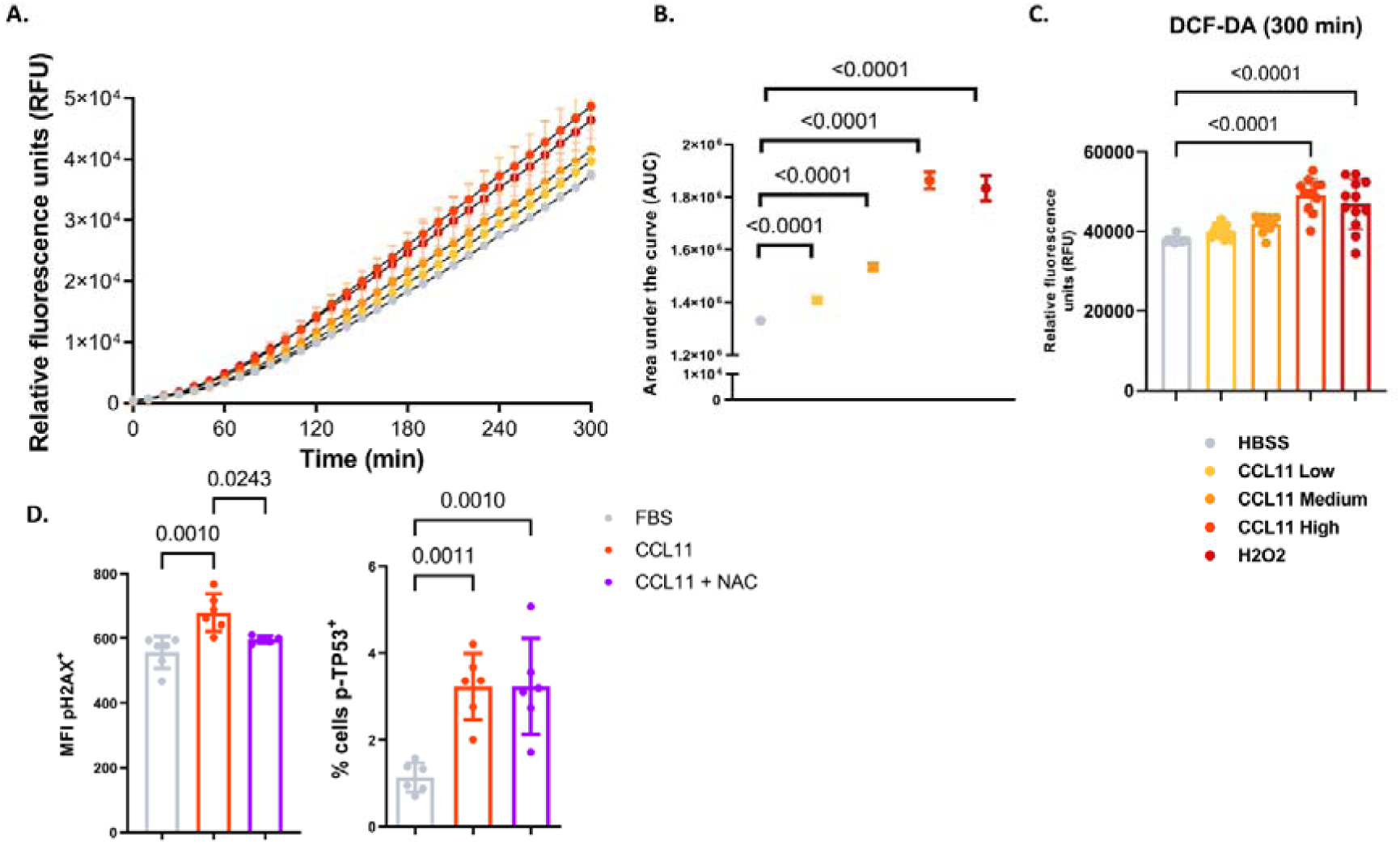
Increasing concentrations of recombinant CCL11 induces ROS formation and DNA damage in PBMC. PBMC were cultured (5x104 cells/well) in 96 well plate and treated with increasing concentrations of rhCCL11 (low: 50ng/mL, medium: 125ng/mL and high: 250ng/mL) **A.** Time course of ROS production quantified via DCF-DA assay after stimulation with increasing concentrations of CCL11. Cells were analyzed using total fluorescence assay in the 96-well plate format at 485/535 nm for 300 minutes. **B.** Absolute ROS quantification from area under the curve (AUC) analysis. **C.** Relative fluorescence (End point, 300 min) of ROS production from DCF-DA assay. **D.** Phosflow analysis showing significant increase in H2AX phosphorylation (pS139) after stimulation with CCL11 (500ng/mL), suppressed by co-treatment with NAC (p-value<0.01). **E.** Phosflow analysis showing significant increase in TP53 phosphorylation (pS37) after stimulation with CCL11 recombinant (500ng/mL) (p-value<0.01). MFI: median fluorescence intensity; NAC: N-Acetyl Cysteine.

## AVAILABILITY

PseudoCell is a free open source regulatory network available in PseudoCell website (http://isenesc.com.br/pages/tools.php) or in the GitHub repository (https://github.com/viniciuspierdona/pseudocell).

